# Metagenome-scale Modeling to Assess Microbiome Metabolic Complementarity for Precision Microbiota Transplantation Therapies

**DOI:** 10.64898/2026.05.15.725570

**Authors:** Zetian Zhang, Mark Holton, Daniel M. Ferrer, Arielle D. Tripp, Alexander Richter, Purushottam D. Dixit, Guillaume Urtecho

## Abstract

Fecal microbiota transplantation (FMT) holds therapeutic promise beyond recurrent *Clostridioides difficile* infection, but clinical outcomes remain unpredictable, in part because existing computational models do not fully capture the metabolic compatibility between donor and recipient communities. Here, we present a metagenome-scale metabolic modeling framework that quantifies metabolic niche complementarity between donor and recipient microbiomes to predict transplantation outcomes. Using MICOM-derived community metabolic models, we show that donor taxa whose metabolic flux profiles are more dissimilar from the recipient community engraft at significantly higher rates in both murine and human FMT cohorts. In a human IBS trial, metabolic models accurately predicted post-FMT community composition via leave-one-out cross-validation and recapitulated disease-associated alterations in short-chain fatty acid, sulfur, and gas metabolism. We then performed 2,548 *in silico* FMT simulations between IBS-D/M patients and donors from the OpenBiome biobank to demonstrate a platform for personalized donor screening. This screen identified super-donors characterized by high taxonomic diversity, broad metabolic niche coverage, and community interaction networks dominated by cross-feeding rather than competition, as quantified by a flux-derived ecological network balance index that strongly predicted engraftment potential. This framework provides a mechanistic, scalable tool for rational donor-recipient matching that could guide personalized microbiome-based therapies.

## INTRODUCTION

The human gut microbiome is a cornerstone of host health, and disruptions in this complex community “dysbiosis” are linked to numerous diseases^1^. Fecal microbiota transplantation (FMT), a procedure in which microbial stool communities from a healthy donor are transferred to a patient, has emerged as a powerful method to restore microbial balance. In recurrent *Clostridioides difficile* infection (CDI), FMT can achieve cure rates around 80-90%, and recently the first standardized FMT products gained FDA approval for prevention of recurrent CDI^2^. Beyond CDI, FMT has been explored as a remedy for metabolic and inflammatory conditions (e.g. obesity, diabetes, IBS), with some reports of symptom improvements^3,4^. However, these broader applications remain experimental, and FMT is not yet routinely used outside of CDI due to variable patient responses and an incomplete understanding of engraftment dynamics^5^.

Despite FMT’s promise, outcomes can differ markedly between recipients. Many patients fail to sustain engraftment of donor bacteria, in part due to colonization resistance, where the native microbiota outcompetes incoming strains for nutrient niches^6^. This niche competition between donor and recipient taxa is thought to be a primary barrier to successful engraftment and successful clinical outcomes^5^. Several groups have attempted to predict FMT success a priori using microbiome sequencing data and machine-learning models^7–10^. For example, Smillie et al. modeled engraftment at the level of individual species and strains, showing that whether a given taxon engrafts can be predicted largely from its donor and recipient abundances and how closely related it is to other bacteria in the community^7^. Other models have taken more holistic perspectives focused on donor-recipient complementarity at community and strain-population levels, and concluded that overall dissimilarity between donor and recipient microbiomes is a dominant driver of FMT colonization patterns rather than properties of any single taxon^10,11^. This concept is based on niche availability and assumes that more dissimilar communities are better able to exploit unfilled niches in the recipient. However, given substantial strain-level variation, taxonomic identity alone is insufficient to assign microbes to their metabolic niches. As a result, we still lack practical, mechanistically-grounded methods to measure functional compatibility between donor microbes and a recipient’s unfilled niches and to use this information prospectively for donor selection.

Ecologically, the gut can be viewed as a mosaic of distinct niches defined by their biochemical resources, spatial locales, and host factors^12,13^. Diverse native microbiota occupy most of these niches, thereby resisting invasion by foreign microbes through competition for nutrients and space^6^. Thus, new microbes can only become established if they are able to exploit a niche that is underutilized by the resident community^14^. Expanding a microbe’s niche range to target resources not consumed by the host’s microbiome has indeed been shown to enable its stable long-term colonization^15^. While identifying niches within microbiomes is challenging, multi-omics approaches now provide direct insight into such niche dynamics. For example, integrated metatranscriptomic and metatranslatomic analyses have classified gut bacteria into functional guilds and identified their preferred nutrient substrates (niches), enabling selective manipulation of community members^16^. While powerful, these empirical approaches are costly, labor-intensive, and they cannot easily capture the full metabolic complexity of natural gut communities. There is a pressing need for practical, scalable methods to profile a patient’s microbiome niche structure so that FMT donors can be rationally chosen to fill vacant metabolic niches.

Genome-scale metabolic models (GEMs) offer a promising computational avenue to characterize microbial niches *in silico*. GEMs leverage genomic annotations and biochemical reaction networks to simulate an organism’s metabolic capabilities, thereby predicting which nutrients it can consume and what metabolites it produces^17^. Recent advances have enabled these models to be scaled up from single strains to whole communities, allowing prediction of microbial activity in complex environments like the gut^18,19^. For instance, the MICOM framework can integrate dozens of gut bacterial GEMs to simulate community metabolism and growth under defined dietary conditions^18^. Such community metabolic modeling has already shown that certain gut bacteria maintain conserved “niche structures” across individuals, and it can predict community-level outputs (e.g. short-chain fatty acid production) in different hosts^20^. Moreover, metabolic modeling has been applied to ecological niche theory, where by treating a microbe’s simulated metabolic phenotype as a proxy for its niche, one can quantify niche overlap or complementarity between species^17^. This approach enables direct comparisons of metabolic niches and provides a quantitative way to partition niches among community members. Recently, these models were used to predict pathogen niches and assess the susceptibility of populations to *Clostridiodes difficile*^*21*^.

In this work, we leverage metagenome-scale community metabolic modeling to develop a metric, metabolic distance, that quantifies niche complementarity between FMT recipients and donor microbes. We show that metabolic distance can accurately capture metabolic patterns that contribute to colonization outcome by building predictive modeling to model engraftment outcome and therapeutic response. We first validate this framework in a controlled murine FMT system without antibiotic pretreatment, demonstrating that donor strains with metabolic capabilities orthogonal to the recipient’s microbiome achieve significantly higher colonization success. We then extend our analysis to a human clinical trial of FMT for irritable bowel syndrome, where no antibiotics were used, to evaluate the model’s predictive power in a translational setting. We then extend this framework into an *in silico* pipeline that performs 2,548 donor-recipient FMT simulations as proof of concept for a clinical tool and identify mechanistic features of super-donors. Our results show that a minimal set of metabolic parameters, measured before transplantation, can accurately predict FMT engraftment outcomes and changes to predicted metabolomes associated with therapeutic response. By integrating metabolic niche profiling with machine learning, this study lays the groundwork for a new approach for matching FMT donors to patients based on complementary microbiome metabolisms to maximize engraftment and therapeutic benefit.

## RESULTS

### Metabolic distance robustly predicts engraftment in a murine FMT model

We first asked whether genome-scale metabolic models (GEMs) could capture quantitative differences in metabolic niche use that explain which taxa engraft during transplantation. GEMs leverage genomic data to model metabolic phenotypes, predicting each microbe’s functional capabilities and ecological role based on simulated nutrient uptake and metabolic outputs under steady-state conditions. We hypothesized that if microbiomes occupy distinct regions of metabolic niche space, then the degree of functional dissimilarity between donor and recipient taxa should predict colonization success, with donor microbes that exploit underutilized niches in the recipient community having a competitive advantage during engraftment.

We tested this hypothesis using a previously published murine FMT dataset^22^. In this work, FMT was performed between all possible pairs of C57BL/6 mice from four different vendors (Jackson Laboratories, Taconic, Envigo, and Charles River), which carried distinct microbiomes, and metagenomic sequencing was used to assess the composition of these communities before and after FMT. Reanalysis reconfirmed substantial variation in engraftment across donor-recipient pairings, with Envigo microbiota transferring the most taxa across all recipients while Taconic communities engrafted the fewest (**Figure S1**). From the assembled metagenome-assembled genomes (MAGs), we used CarveME to reconstruct GEMs and applied MICOM to simulate steady-state metabolic fluxes under a high-fiber diet.

We next examined whether the four vendor microbiomes differed in their coverage of metabolic niche space. We performed PCoA on MICOM-derived exchange flux profiles using cosine distance, which compares the proportional distribution of fluxes across metabolites and is therefore sensitive to differences in metabolic strategy independent of absolute flux magnitude.

Microbial taxa clustered largely according to their taxonomic order, yet we also observed substantial variability within each order indicating that closely related microbes can have markedly different metabolic potentials (**Figure 1A**) ^23^. We then divided the resulting two-dimensional niche space into a 20 × 20 grid to evaluate cell occupancy across microbiomes. The Envigo microbiome occupied the greatest number of grid cells (n=67), followed by Charles River (n=56), Jackson Laboratories (n=45), and Taconic (n=35) (**Figure 1B**). This ranking corresponded to prior observations of engraftment potential in this system, where Envigo was the most successful donor and Taconic the least^22^, suggesting that broader metabolic niche coverage may confer a colonization advantage.

**Figure 1.**
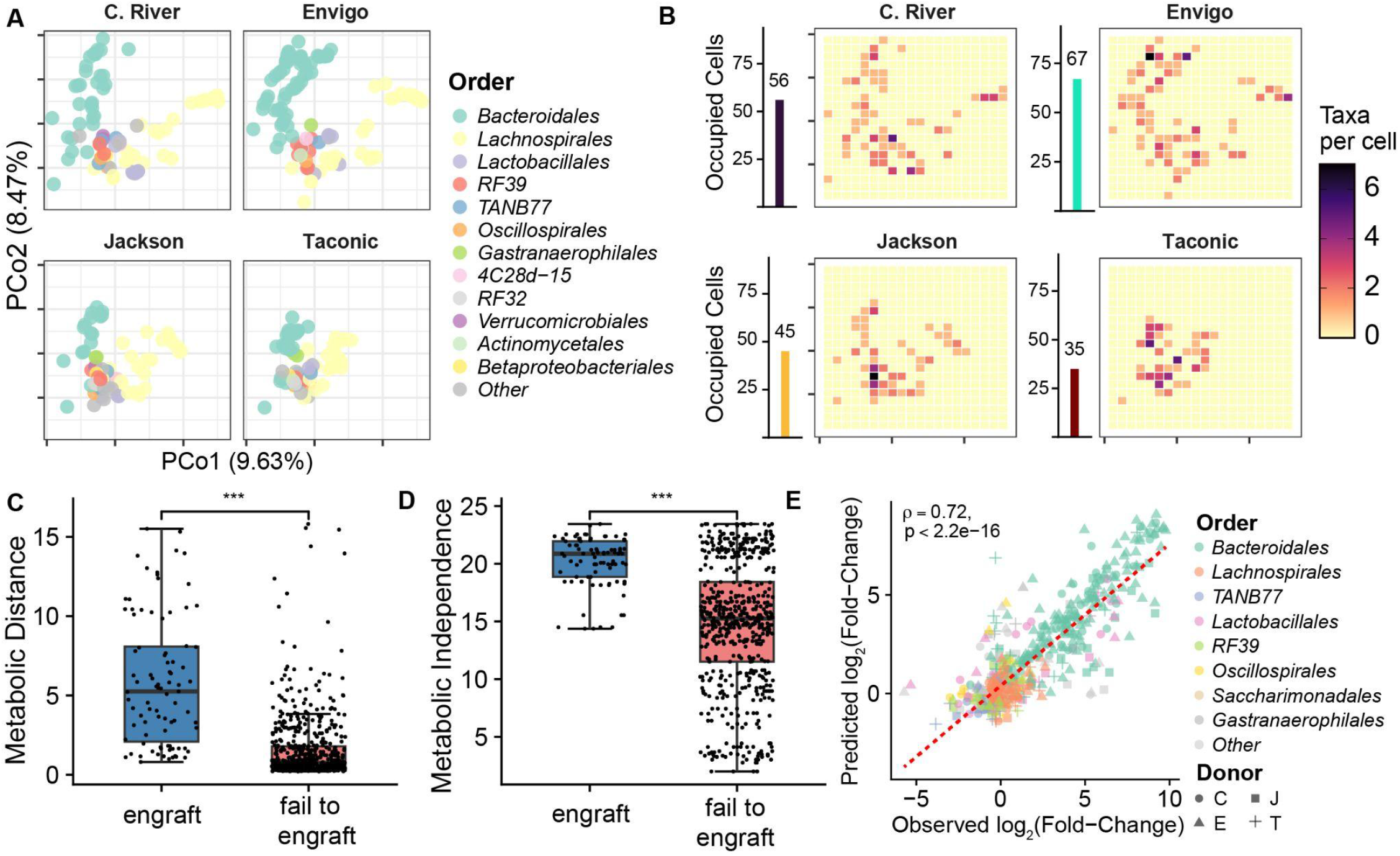
Metabolic distance predicts engraftment outcomes in a murine FMT model. **A)** Principal coordinates analysis (PCoA) of MICOM-derived metabolic exchange flux profiles for metagenome-assembled genomes (MAGs) from four vendor-distinct murine microbiomes (Charles River, Envigo, Jackson, Taconic). Each point represents one MAG, colored by taxonomic order. **B)** Grid occupancy of metabolic niche space. A shared 20 × 20 grid was overlaid on PCoA coordinates across all four communities; tile color indicates the number of taxa per cell. Inset bar plots show the total number of occupied grid cells for each community. **C)** Metabolic distance of donor taxa to the recipient community for taxa that successfully engrafted versus those that failed to engraft (p < 2.2 × 10^−16^, two-sided Mann-Whitney U test). **D)** Metabolic independence scores for engrafting versus non-engrafting taxa (p < 2.2 × 10^−16^, two-sided Mann-Whitney U test). **E)** Random forest regression predicting post-FMT changes in MAG relative abundance. Each point represents one MAG across all donor-recipient pairings, colored by taxonomic order and shaped by donor identity. Dashed line indicates the linear fit. Predictions shown are held-out values from 10-fold cross-validation.

To formalize this relationship, we developed a quantitative metric of metabolic niche complementarity that could be applied directly to FMT. We defined “metabolic distance” as a measure that compares the steady-state metabolic flux profiles of microbes across donor and recipient microbiomes (see Methods). This metric uses simulated fluxes to compute pairwise distances between microbes based on the degree of overlap or complementarity in their metabolic outputs and nutrient use. For each pairwise donor-recipient cohort, we calculated metabolic distance between all MAGs present in the corresponding microbiomes. In parallel, we used Anvi’o to estimate metabolic independence for each MAG, a metric that evaluates the comprehensiveness and completeness of metabolic pathways within genomes, where microbes with more complete pathways are considered more independent. Comparing these values to FMT outcomes, we found that taxa that successfully colonized recipients exhibited significantly greater metabolic distance from the recipient community (**Figure 1C**, p < 2.2 × 10^−16^, Mann-Whitney U test). We evaluated several alternative distance metrics for this comparison, including L1, L2, and cosine distance, all of which separated colonizers from non-colonizers in the same direction but with less effect size than Mahalanobis distance (**Figure S2**). Successfully colonizing taxa were also more metabolically independent (**Figure 1D**, p < 2.2 × 10^−16^, Mann-Whitney U test) than non-engrafting taxa.

Given that metabolic distance and metabolic independence were strongly associated with engraftment, we next evaluated whether these features, together with simple community descriptors, could predict colonization outcomes. We trained a random-forest classifier to predict colonization outcomes using metabolic distance, metabolic independence, the pre-FMT abundances of MAGs in donor and recipient, and the ratio of Shannon diversity scores between donor and recipient; which has been previously associated with greater transfer during FMT^22^. This classifier accurately separated colonizing from non-colonizing taxa across all donor-recipient pairings (auROC = 0.998; prAUC = 0.795; null prAUC = 0.148 **Figures S3A, S3B**) and maintained a median auROC of 0.918 even when trained on only 10 percent of the data (**Figure S3C**). We used ANOVA to assess the amount of variance explained by each parameter and found that metabolic distance was the strongest predictor of colonization, explaining 41.5% of the variance in colonization outcomes (**Figure S3D**). Using the same model architecture, we trained a random forest regression model to quantitatively predict changes in MAG relative abundance within FMT recipients. This model predicted post-FMT changes in abundance with an average Spearman rho of 0.718 ± 0.045 accuracy across 10-fold cross-validation (**Figure 1E**, p < 2.2 × 10^−16^). Collectively, these findings show that metabolic distance, derived from community-scale metabolic models, provides a simple yet powerful predictor of which microbes will engraft during FMT and the resulting composition of recipient communities.

### Metagenomic and Metabolic Signatures of IBS-D/M

Irritable bowel syndrome with diarrhea or mixed stool pattern (IBS-D/M) is a prevalent functional gastrointestinal disorder that imposes substantial costs on patients and health systems. Multiple studies implicate the gut microbiome in IBS pathophysiology, including alterations in community composition, gas production, and fermentation profiles. Microbiome-targeted interventions such as FMT and defined microbial therapeutics have shown promise in subsets of patients, but clinical responses are highly variable, and the microbial or metabolic features that distinguish responders from non-responders remain poorly defined.

We reasoned that metabolic modeling could capture key microbial and metabolic signatures that shed light on IBS pathophysiology and discriminate clinical responders from non-responders to FMT. To test this, we performed a retrospective analysis of a study that assessed the efficacy of FMT in treating IBS-D/M patients^24^. This study implemented a randomized, double-blind, placebo-controlled, parallel-group, single-center study (n=30) for IBS. Participants met ROME III criteria for IBS-D or IBS-M, ranged from 18 to 75 years old, and exhibited moderate to severe symptoms based on the IBS Severity Scoring System (IBS-SSS ≥175). Stool samples were collected for metagenomics sequencing from donors, and from participants at baseline (T0), 6-month (T6) and 12-month (T12) follow-ups. At 3-months, patients were reassessed for IBS and deemed ‘responders’ if they achieved a 75 score decrease in IBS-SSS^25^. Importantly, no antibiotic pretreatment was administered to study participants prior to FMT, making this dataset a comparable extension of our murine FMT model.

Microbiome analysis revealed significant compositional differences between healthy donors and IBS patients before FMT, including a significant increase in *Bacteroidales* species in donors (**Figure S4**). To link these patterns to clinical response, patient microbiomes at T6 were compared to their respective T0 baselines, with differences evaluated separately for responders and non-responders while accounting for patient-specific variation across time points. Responders showed strong enrichment of *Bacteroidales* following FMT, whereas non-responders exhibited minimal compositional change, indicating a positive association between transfer of *Bacteroidales* and remission of IBS symptoms (**Figure 2A**).

**Figure 2.**
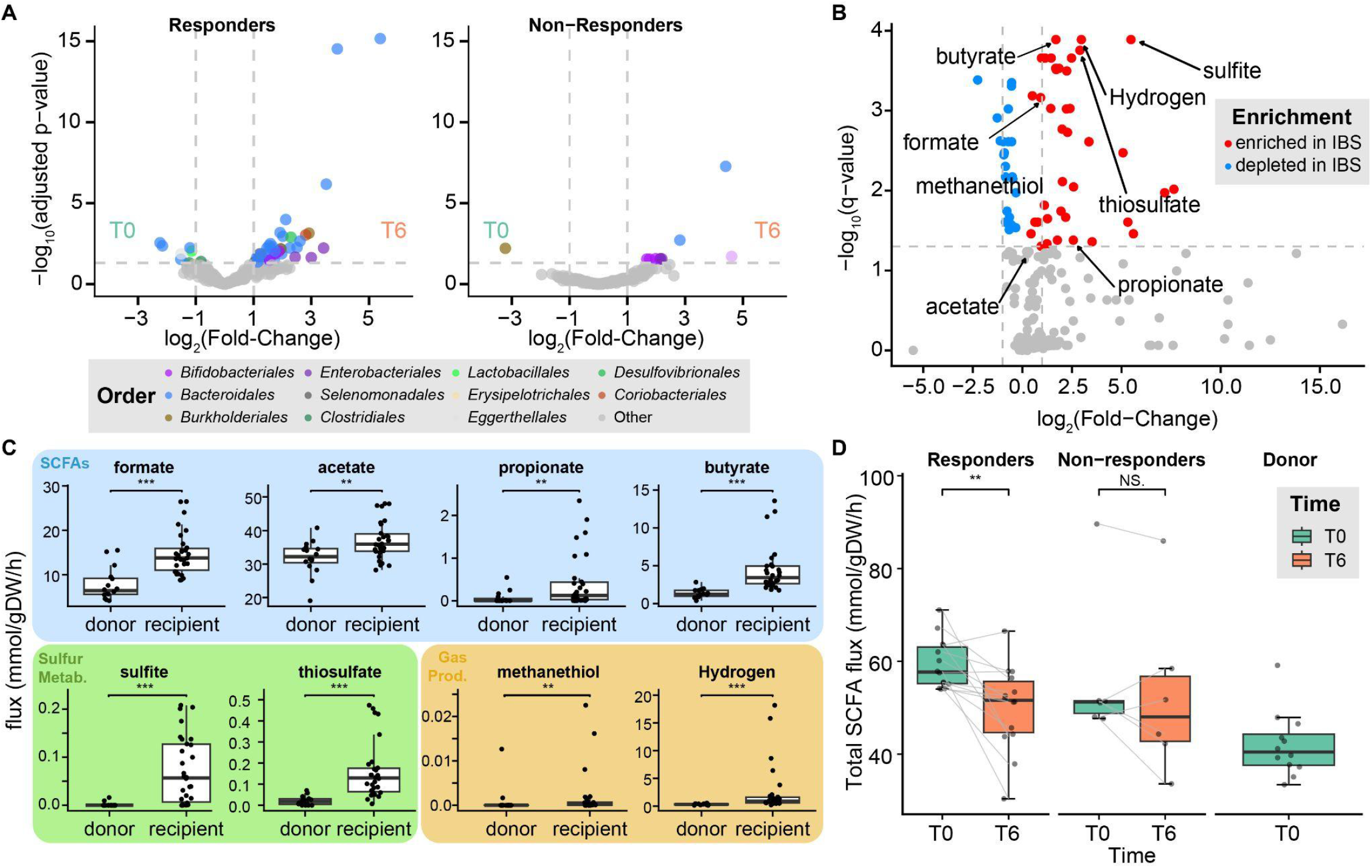
Community-scale metabolic models recapitulate compositional and metabolic signatures of IBS-D/M. **A)** Differential abundance analysis comparing overall recipient microbiome composition between baseline (T0) and six months post-FMT (T6) for responders (left) and non-responders (right). Each point represents one taxon, colored by taxonomic order. The x-axis shows log2-normalized fold change (T6 vs T0) and the y-axis shows -log10 adjusted p-value. Responders showed significant enrichment of Bacteroidales at T6, while non-responders exhibited minimal compositional change. **B)** Differential metabolic flux analysis between IBS patients at baseline and healthy donors, computed from MICOM community-scale metabolic models. Each point represents one secreted metabolite; red indicates metabolites with higher predicted production in IBS patients and blue indicates metabolites with higher predicted production in donors. Key metabolites are labeled, including SCFAs (butyrate, formate, propionate, acetate), sulfur metabolites (sulfite, thiosulfate), and gas products (hydrogen, methanethiol). Dashed line indicates significance threshold. **C)** Predicted community export fluxes for individual SCFAs (blue, top row), sulfur metabolites (green, bottom left), and gas products (yellow, bottom right) in donors versus IBS recipients at baseline. IBS patients showed significantly elevated production across all metabolite classes. Boxplots show median and interquartile range with individual data points overlaid; significance determined by two-sided Mann-Whitney U test (**p < 0.01, ***p < 0.001). **D)** Total predicted SCFA flux (sum of formate, acetate, propionate, and butyrate) at T0 and T6 for responders, non-responders, and donors. Lines connect paired samples from the same patient. Responders showed a significant decrease in total SCFA production following FMT (p < 0.01), whereas non-responders showed no significant change (NS).

We then explored whether differences in microbial composition also drove differences in metabolic activities of the microbiome that could indicate disease. Specifically, we evaluated whether community-level metabolic models could capture these differences and provide insight into molecular drivers of IBS-D/M development and remission. We constructed community metabolic models by mapping metagenomic reads from the Goll et al. IBS cohort to genomes from the AGORA database ^26^, a set of 818 genomes from human gut microbiomes with curated GEMs. After generating models for all donors and recipients at T0 and T6, for each of the 468 molecular products secreted by each community, we calculated a weighted export flux, which was defined as the sum of the product of the normalized export flux and the relative abundance of each microbe. IBS patient microbiomes showed significantly higher predicted production of short-chain fatty acids (SCFAs), sulfur metabolites, and gas (e.g. hydrogen and methane) prior to FMT compared to healthy donors (**Figure 2B, 2C**). These computational predictions are consistent with clinical reports of altered SCFA profiles, gas production, and sulfide production in IBS patients^27,28^. Thus, these community-scale models capture metabolic signatures characteristic of IBS-D/M.

Given that the predicted alterations in key metabolic properties in the IBS cohort by our metabolic models were consistent with clinical data, we next focused on whether SCFA productions can serve as a key metabolic readout for remission of IBS following FMT. Using weighted export flux, we defined total SCFA production as the sum of the total flux of propionate, acetate, butyrate, and formate and compared it between T0 and T6 for this IBS cohort. Responders showed a significant decrease in total SCFA production at T6 relative to their T0 baseline, reaching levels comparable to those of donors (**Figure 2D**). However, there were no significant overall differences in total SCFA flux between responders and non-responders at T6, suggesting that additional factors, such as non-SCFA metabolites, may drive symptoms in these non-responders to FMT. Together, these results indicate that metabolic modeling captures key compositional and functional features of IBS-D/M and that successful response to FMT is associated with engraftment of *Bacteroidales* and a reduction in community SCFA output.

### Systematic donor-recipient mapping for predictable FMT outcomes

Building on our findings that metabolic models predict engraftment and capture IBS-D/M-associated metabolic signatures, we next asked whether these metagenome-scale models could be used prospectively to guide donor selection for FMT. Specifically, we sought to test whether the same modeling framework could be repurposed to identify donor-recipient combinations that maximize engraftment and drive metabolomic shifts associated with clinical response.

We first assessed whether our metabolic modeling approach could accurately predict FMT outcomes in human IBS studies. Using metagenome-scale metabolic models constructed from IBS-D/M patients prior to FMT, we computed metabolic distance between patients and their actual donors and predicted the composition of patients six months after treatment. Predicted and observed post-FMT compositions were strongly concordant across leave-one-recipient-out cross-validation (Spearman rho = 0.76 ± 0.14, p < 2.2 × 10^−16^; **Figure 3A**), and prediction accuracy was consistent for both responders and non-responders (**Figure 3B**). To examine the association between engraftment and remission, we calculated engraftment scores by measuring the Bray-Curtis distance of each patient’s post-FMT microbiome relative to their pre-FMT baseline and their donor. By this metric, clinical responders shifted significantly closer to their donors than non-responders (p < 0.05, Wilcoxon rank-sum test; **Figure 3C**), indicating that greater donor-like engraftment is associated with symptom improvement. Together, these results demonstrate that metabolic distance-based models generalize from murine to human FMT cohorts and that the degree of predicted engraftment tracks with clinical outcomes.

**Figure 3.**
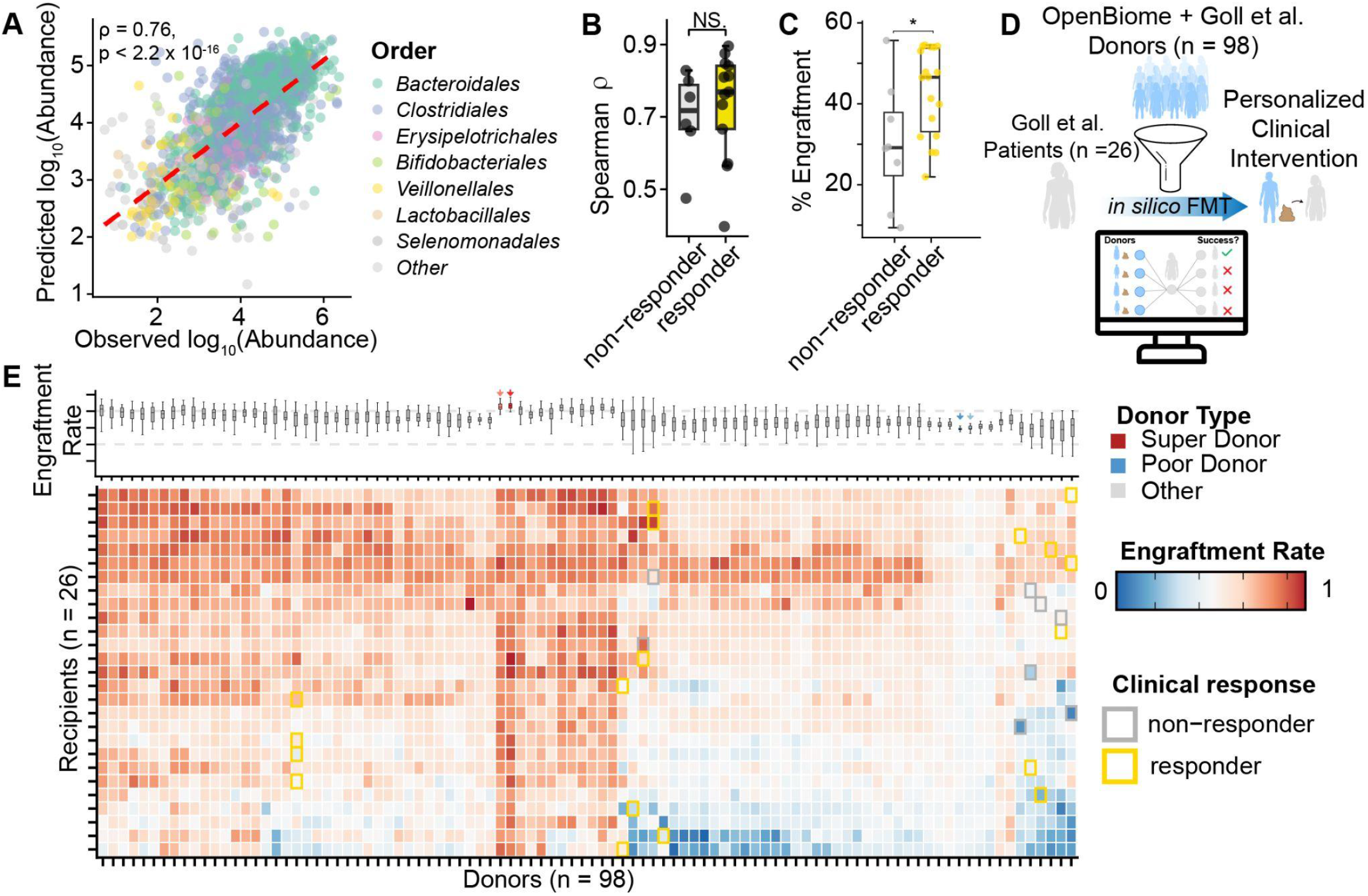
Metabolic models predict post-FMT composition in human IBS and enable *in silico* donor screening. **A)** Leave-one-recipient-out cross-validation of a random forest regression model predicting post-FMT taxon abundance from metabolic distance, metabolic independence, pre-FMT abundances, and donor-to-recipient diversity ratio. Each point represents one taxon for one recipient, colored by taxonomic order. The red dashed line indicates the linear fit; Spearman rho and p-value are annotated. **B)** Per-recipient Spearman rho between predicted and observed post-FMT abundance, stratified by clinical response. Each point represents one recipient held out during model training. NS, not significant (Wilcoxon rank-sum test). **C)** Engraftment scores for responders and non-responders, calculated as the relative shift in Bray-Curtis distance from the pre-FMT composition toward the donor composition at six months post-FMT (*p < 0.05, Wilcoxon rank-sum test). **D)** Schematic of the *in silico* FMT screening framework. Metabolic models from 26 Goll et al. IBS patients, 12 Goll et all donors, and 86 OpenBiome donors were used to simulate 2,548 donor-recipient pairings and predict engraftment outcomes. **E)** Heatmap of predicted engraftment rates across all donor-recipient combinations. Rows represent recipients (n = 26) and columns represent donors (n = 98), ordered by hierarchical clustering. Color intensity reflects predicted engraftment rate (red = high, blue = low). Bordered tiles indicate actual donor-recipient pairings from the Goll et al. trial (gold = responder, gray = non-responder). Boxplots above show the distribution of predicted engraftment across recipients for each donor; arrows indicate super-donors (red) and poor donors (blue).

We then asked whether this framework could serve as a practical platform to screen and select donors for personalized FMT. We used the same modeling framework to perform *in silico* FMT simulations for 2,548 independent FMTs between all 26 IBS-D/M patients from this study and 98 candidate donors, including 12 from the original Goll et al. cohort and 86 individuals characterized by the OpenBiome project (**Figure 3D**). This analysis revealed substantial heterogeneity in predicted engraftment across donor-recipient pairs, driven by both donor and recipient identity (**Figure 3E**). To determine whether donor or recipient identity is the primary driver of FMT engraftment, we subsampled 50% of all simulated donor-recipient combinations across 100 iterations and calculated the proportion of total variance in engraftment explained by each factor. Recipient identity explained a median of 41.7% of variance compared to 34.9% for donor identity, indicating that recipient-specific features play a larger role in determining engraftment outcomes (Wilcoxon rank sum test, p < 2.2 × 10^−16^). Current approaches to FMT donor selection focus on identifying universal “super-donors” while remaining largely agnostic to recipient characteristics, yet these results suggest that matching donors to recipients based on recipient-specific features may be more effective than optimizing donor selection alone.

### Community characteristics of ‘Super Donor’ Microbiota

While the existence of ‘super donors’ in FMT has been suggested^29^, the community ecological characteristics have not been deeply investigated. While we found that recipient identity largely determined FMT outcomes, two donors appeared to be the exception to this, exhibiting consistently high predicted engraftment across nearly all simulated recipients. These donors, henceforth referred to as “super-donors” 1 and 2, showed robust engraftment with a median engraftment score of 0.83 ± 0.064 and 0.81 ± 0.059. To shed more light on the community ecological characteristics of super-donor microbial communities, we performed a series of community analyses. The super-donors harbored some of the highest Shannon diversity values among all donors, which strongly correlated with engraftment outcomes across the cohort (**Figure 4A**, R^2^ = 0.793, p < 2.2 × 10^−16^, GAM). Moreover, taxa from super-donors were, on average, more metabolically distant from recipients than taxa from other donors (**Figure 4B**, p < 2.2 × 10^−16^, Mann-Whitney U test), consistent with our model that complementary, orthogonal metabolisms promote engraftment. Analyzing the distribution of metabolic independence scores within these communities, the two best donors were less metabolically independent on average (p < 2.2 × 10^−16^, Mann-Whitney U test; **Figure 4C**), with differences largely attributable to a bimodal distribution in metabolic independence scores that was absent in poorly-engrafting donors. This bimodal distribution suggested a community composed of a mixture of metabolically self-sufficient and cross-feeding-dependent taxa, implying extensive within-community metabolic cooperation.

**Figure 4.**
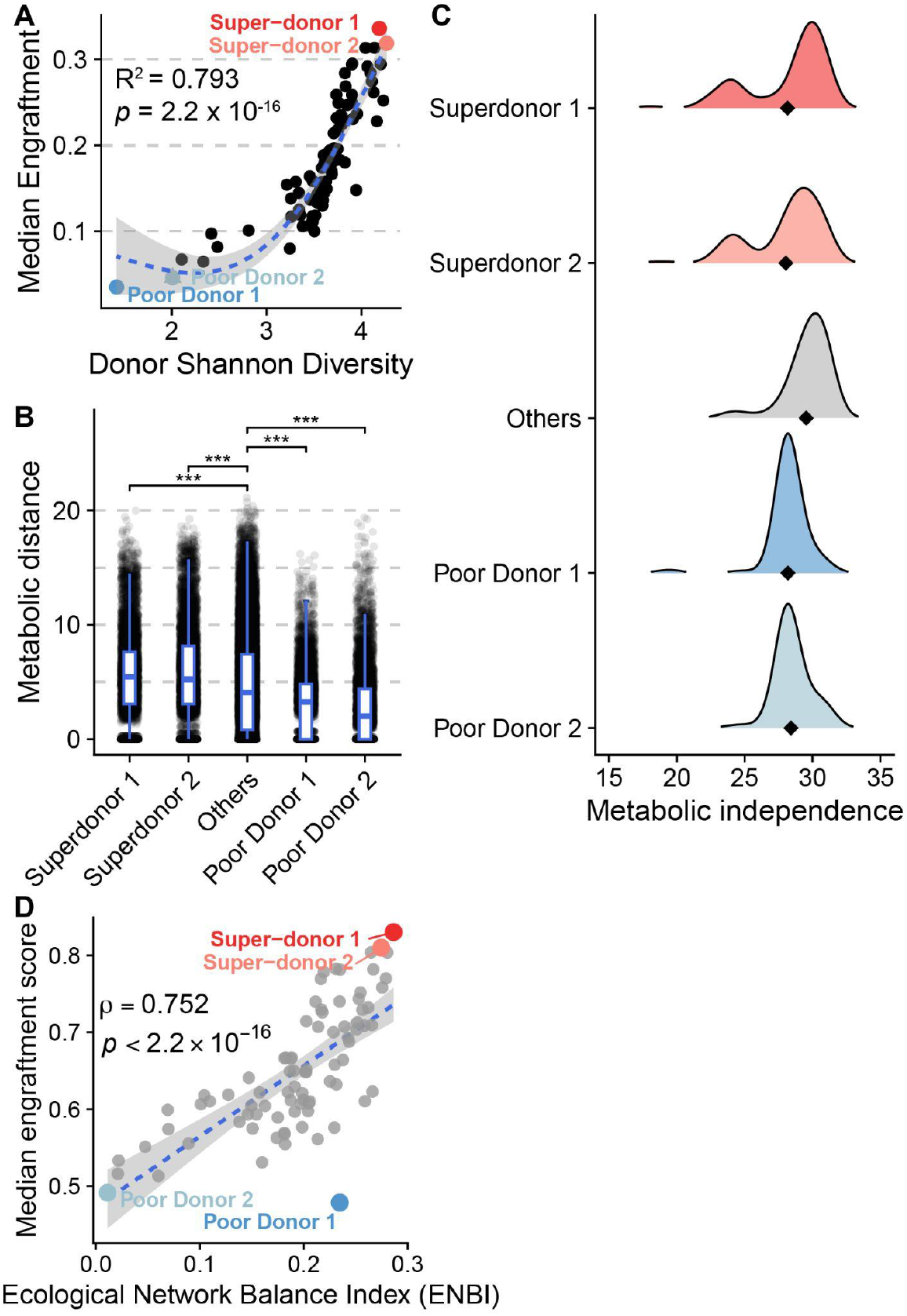
Community ecological characteristics of super-donors. **A)** Relationship between donor Shannon diversity and median predicted engraftment across all recipients. Each point represents one donor; the blue line shows a generalized additive model (GAM) fit with 95% confidence interval (R^2^ = 0.793, p < 2.2 × 10^−16^). Super-donors 1 and 2 (red) occupy the high-diversity, high-engraftment extreme; poor donors 1 and 2 (blue) cluster at the low end. **B)** Distribution of metabolic distance to recipients for taxa from super-donors, other donors, and poor donors. Super-donor taxa exhibited significantly greater metabolic distance from recipient communities than taxa from other donors (***p < 0.001, Mann-Whitney U test). Boxplots show median and interquartile range with individual data points overlaid. **C)** Density distributions of metabolic independence scores for taxa within each donor category. Diamonds indicate the median. Super-donor communities displayed a bimodal distribution of metabolic independence, with a subpopulation of low-independence taxa absent in poor donors. **D)** Relationship between the ecological network balance index (ENBI), computed from MICOM-derived pairwise metabolic fluxes, and median engraftment score across all donors. Each point represents one donor; with super-donors (red) and poor donors (blue)labeled. The ENBI quantifies the relative dominance of positive (cross-feeding) over negative (competitive) interactions within each donor community (Spearman rho = 0.752, p < 2.2 × 10^−16^). Dashed line shows the linear fit with 95% confidence interval.

The reduced average metabolic independence of super-donor communities implies that their taxa are substantially reliant on cross-feeding to meet their metabolic demands. Corral Lopez et al. (2026)^30^ showed that tightly cross-feeding consortia form compact metabolic loops that efficiently exploit available resources, allowing them to outcompete and displace resident taxa. We reasoned this dynamic may similarly operate in the FMT context, where super-donor consortia with analogous cross-feeding architectures could more effectively overcome colonization resistance in the recipient gut. Inspired by the ecological network balance index (ENBI) by Corral Lopez et al. (2026), which quantifies the relative balance of positive and negative interactions within a community, we computed an analogous metric using MICOM-derived metabolic fluxes to evaluate pairwise competitive and cross-feeding interactions among all donor community members (see Methods). This analysis showed that flux-derived interaction balance correlated strongly with engraftment (**Figure 4D**, ρ = 0.752, p < 2.2 × 10^−16^), suggesting that the cross-feeding network architecture characteristic of super-donor communities is a key determinant of engraftment potential.

## DISCUSSION

In this study, we used metagenome-scale metabolic modeling to apply ecological theory of niche complementarity towards rational donor selection in FMT. By leveraging MICOM-based community models and a new metric of metabolic distance, we showed that donor taxa that are more metabolically dissimilar from the recipient microbiome are more likely to engraft in a murine FMT model. Metabolic distance and metabolic independence together explained a substantial fraction of engraftment variance and enabled accurate prediction of colonization outcomes using simple machine learning models. We then extended this framework to a human IBS-D/M FMT trial, where community-scale models recapitulated disease-associated metabolic signatures and distinguished responders from non-responders based on predicted fluxes. Finally, we demonstrate an approach to performing 2,548 *in silico* FMT screens across all donor-recipient combinations, using model-derived features to predict engraftment rates for each pairing and thereby map how donor choice determines post-FMT states. Across all donors, we identified two super-donors whose taxa occupied a broader region of metabolic niche space, achieved consistently high predicted engraftment across diverse recipients, and provided a functional explanation for the super-donor phenomenon. Together, these results support a mechanistic view in which metabolic niche complementarity, rather than taxonomic similarity alone, drives FMT engraftment and the resulting shifts in microbiome function.

Our findings build upon existing models for predicting FMT success that have focused primarily on taxonomic or pathway composition, phylogenetic relatedness, or simple diversity metrics^7,8,10^. Prior work has shown that donor-recipient similarity in community structure or the presence of particular taxa can correlate with FMT outcomes, yet these features do not directly capture how communities use resources or partition metabolic functions^10^. Here, we show that a single function-derived feature, metabolic distance, is sufficient to accurately predict which taxa will colonize and their relative abundances after transplantation. This is consistent with ecological theory in which successful invaders occupy underutilized niches and face less direct competition from resident species^14,15^. By embedding high-dimensional flux profiles into an interpretable niche space, our framework moves beyond black-box correlational models and provides a mechanistic link between metabolic complementarity and colonization resistance.

These insights provide guiding principles for FMT donor selection. The two super-donors identified in our i*n silico* screen were characterized by high taxonomic diversity, broad metabolic niche coverage, and a bimodal distribution of metabolic independence scores, reflecting communities composed of diverse metabolisms consisting of both self-sufficient taxa and cross-feeding-dependent specialists. Consistently, a flux-derived analog of the ENBI^30^ scored highest in super-donors, suggesting that cross-feeding-dominated interaction networks confer a colonization advantage by enabling tightly cooperative consortia to exploit underutilized metabolic niches in the recipient gut. Notably, Corral López et al. found that elevated ENBI values characterize dysbiotic rather than healthy communities, where tightly cross-feeding consortia outcompete resident taxa and dominate the community. Our results suggest that this same competitive mechanism may operate during FMT, where incoming donor consortia with strong internal cross-feeding networks can similarly overcome colonization resistance in the recipient. These findings suggest that donor screening should move beyond diversity and taxonomic criteria to explicitly quantify metabolic niche coverage and interaction network balance, metrics that can be computed prospectively from metagenomic data. Importantly, because recipient identity explained more variance in engraftment than donor identity, effective donor matching will require joint consideration of the donor’s niche profile and the recipient’s existing metabolic landscape.

Our retrospective analysis of an IBS-D/M FMT trial captured known features of IBS, including altered SCFA, sulfur, and gas metabolism, and showed that responders to FMT shifted toward donor-like metabolic states. Although engraftment rate trended with successful response to FMT, several patients with high engraftment did not experience remission of IBS symptoms^11^. This suggests that high engraftment alone is not sufficient, and that donor selection should prioritize microbiomes whose engrafted members actively reduce disease-associated metabolites. With further refinement of environmental priors, metagenome-scale models may be able to predict these effects, enabling FMT strategies that are guided not only by engraftment probability but also by anticipated changes in microbiome metabolic phenotypes. Such an approach could support targeted and personalized care across diverse microbiome-associated diseases.

This work lays a foundation for several future developments. While our models use curated GEMs and metagenomic data, they still rely on steady-state assumptions and incomplete knowledge of strain-level variations in metabolism. Integrating time-resolved data, diet records, and host-specific constraints could improve estimates of engraftment likelihood as well as present opportunities to predict metabolic changes and downstream clinical impact. Applying this framework to rationally designed consortia^19^ would allow direct experimental evaluation of whether engineered niche complementarity can match or surpass the performance of natural donors. Finally, extending these ideas to additional diseases and patient populations beyond IBS-D/M will be essential for establishing metabolic niche complementarity as a general principle for effective microbiome-based treatments.

## METHODS

### Study design and cohorts

Three shotgun metagenomic datasets were analyzed in this study. The first comprised murine fecal samples from C57BL/6 mice sourced from four vendors: Taconic Biosciences, Envigo, Jackson Laboratories, and Charles River that were characterized in a previous study^22^. Eight mice per vendor were used, and cohousing was performed in pairs, with each mouse participating in two pairwise cohousings. Samples collected before and after all pairwise cohousings were included, enabling analysis of both vendor-associated microbiome differences and inter-vendor microbiome transfer dynamics.

The second dataset was derived from a human clinical trial (Goll et al. 2020), in which participants with irritable bowel syndrome (IBS) underwent fecal microbiota transplantation (FMT). From the 83 participants enrolled in the REFIT study, only those in the active treatment arm with complete sample sets were included, yielding 22 participants with stool collected at three time points. Participants were stratified by treatment outcome into responders (n=14) and non-responders (n=8). For the *in silico* FMT simulations, an additional four patients were included, for a total of 26, although these individuals did not return for 6 month and 12 month follow-ups so were excluded from previous analyses.

A third dataset used in this work was from the Openbiome dataset^31^, comprising stool samples from healthy donors screened by OpenBiome, a non-profit stool bank. From the full cohort of 90 subjects sampled, the subset of 86 donors for whom shotgun metagenomic sequencing data were available was used in the present analysis. Donors were healthy adults aged 19-45 years, free of gastrointestinal, autoimmune, cardiovascular, neurological, and metabolic conditions at recruitment, screened for pathogens, and were fully deidentified prior to sample receipt.

### Metagenomic sequencing processing and abundance profiling

Read trimming, adapter removal, and quality filtering were performed using fastp^32^ (v1.0.1) with polyG and polyX tail trimming enabled (-g -x) and a custom adapter FASTA supplied via --adapter_fasta. Taxonomic classification of cleaned reads was performed using Kraken2^33^ (v2.17.1) against the standard 8 GB database, followed by abundance re-estimation at the species level using Bracken; per-sample Bracken reports were subsequently merged into a single species-level count table. Read identifiers classified as belonging to AGORA-relevant taxa were extracted by cross-referencing Kraken2 output against a predefined ancestral taxa lineage list using csvgrep and csvcut, and matching reads were isolated from the cleaned FASTQ files using a custom filtering script. Filtered reads were then aligned to an indexed AGORA genome database using minimap2^34^ in short-read mode (-x sr) with secondary alignments suppressed (--secondary no). Resulting alignments were filtered using samtools (v1.21) to retain only properly paired, first-in-pair reads mapped in the correct orientation (flags -f 0 × 42 -F 0xF0C), and per-taxon read counts were extracted from the alignment via an inline Perl script parsing the taxon ID from the reference sequence name. Final per-sample count files were aggregated into a single count matrix using a custom Python script. All parallelizable steps were distributed across samples using GNU Parallel (v20250422).

### Genome-scale metabolic model reconstruction and community modeling (CarveMe, MICOM)

The mouse data described in this study was acquired from a publicly available dataset, in which the raw metagenomic sequencing data was processed under a standardized pipeline to generate individual contig assemblies, abundance profiles, and taxonomic annotations^22^. Assemblies from this study were further processed by CarveMe v1.6.0^35^ with gapfilling to generate the reference model database GEMs. For each mouse cohort, namely Jackson, Envigo, Taconic, and Charles River, a pre-FMT and a post-FMT model were constructed using MICOM v0.37.0 with “cutoff=0”^18^. Read counts were averaged within each cohort, and taxonomic information was summarized at genus level to match the reference database prior to model construction. Raw metagenomic read counts were normalized by genome length, to account for read counts for a given taxon being proportional to both its cellular abundance and its genome length. Growth was simulated using the AGORA high fiber diet with “tradeoff=0.7” and genus-level pairwise interaction networks were derived from the resulting community growth models.

Abundance and taxonomy data derived from the human clinical IBS FMT treatment data and healthy Openbiome donor data were summarized at species level. Prior to model construction, for each sample, read counts from each species were further normalized by their respective genome lengths to control for bias by uneven sequencing depth. Model construction was performed with “cutoff=0.001”. Predicted metabolic fluxes and growth rates were generated using MICOM’s *grow* function with “tradeoff=0.7” and a standard AGORA western diet for downstream analysis. Species-level pairwise interaction networks were built with MICOM’s interactions function.

### Metabolic feature engineering (metabolic distance, metabolic independence, diversity)

Genus/species-level pairwise interaction data was used to calculate metabolic distance between two samples. Briefly, interaction flux estimates were first aggregated across biological replicates by metabolite and interacting species, producing a mean flux profile that represents the metabolic capacity of each taxa. Based on the predefined pairings between donors and recipients from the FMT treatment, for each comparison both donor- and recipient-specific profiles were extracted from the aggregated dataframe and were reshaped to taxon x metabolite matrices. Mahalanobis distance was used to compute pairwise distances between donor and recipient taxa in metabolite space. Because exchange fluxes are highly correlated across metabolic pathways, Mahalanobis distance accounts for the covariance structure of the metabolite space, preventing correlated fluxes from inflating distance estimates. The covariance matrix was estimated using a shrinkage estimator to ensure numerical stability given the high dimensionality of the flux data relative to the number of taxa. The resulting matrix captures the metabolic dissimilarity between every donor-recipient taxon pair. To summarize each donor taxon’s metabolic distance from the recipient community, we computed the mean distance to its 15 nearest recipient taxa, representing the most likely niche competitors for that organism. Metabolic independence was computed for each AGORA genome using the anvi-script-estimate-metabolic-independence function in Anvio v8.0^36^ and genome score was extracted.

### Engraftment and outcome definitions (colonization, composition shift, clinical response)

Raw abundance counts were first normalized by taxon-specific genome length and then converted to relative abundance within each sample. Only taxa present in at least 5% of the samples with relative abundance exceeding 0.01% were retained. The raw count matrix was then filtered to retain only this subset of taxa for differential abundance analysis, which was performed to distinguish the compositional shift between responders and non-responders for each group between T0 and T6 using DESeq2 with the formula “∼subject_id + time” to control for patient and time effect^37^.

To assess donor microbiome colonization following FMT treatment, an engraftment score was defined as the percent change in Bray-Curtis distance between the recipient microbiome at T6 and T0 relative to the donor microbiome at T0. For this analysis, the abundance profile was normalized by genome length and converted to relative abundance but was not subjected to the filtering criterion described above. Normalization by genome length is necessary as raw metagenomic read counts for a given taxon are proportional to both its cellular abundance and its genome length. The score was further transformed to 1 - percent change to represent engraftment similarity.

Metabolic differential enrichment analysis was performed by first converting MICOM’s flux dataframe into sample-level production rates using MICOM’s production_rates function followed by MICOM’s compare_group function.

### Predictive modeling and model evaluation (classification, regression, CV, ANOVA/importance)

To predict the colonization outcome of individual donor taxa following FMT, a random forest classifier was trained using the ranger package (R) with five metabolic features calculated for each OTU: metabolic distance, metabolic independence, donor taxon relative abundance at baseline (log_10_-transformed), recipient OTU abundance at day 0, and the donor-to-recipient Shannon diversity ratio for the whole microbiome. The classifier was trained with 50 trees and mtry = 2, with colonization status of each OTU was predicted as a binary outcome. Model performance was evaluated using out-of-bag (OOB) AUROC computed via the pROC package, and robustness was further assessed through Monte Carlo cross-validation across 19 iterations at training set proportions ranging from 5% to 90%. Precision-recall performance was evaluated over 20 iterations at a 30/70 train/test split using the PRROC package, with a null model generated by label permutation as a baseline comparator. Feature importance was assessed using the Gini impurity criterion. A complementary random forest regression model was additionally trained to predict the magnitude of colonization as log-transformed fold-change, using 150 trees with permutation-based importance, and was evaluated using OOB R^2^, Pearson correlation, and Spearman rank correlation between observed and OOB-predicted values.

### *In silico* donor-recipient screening and super-donor analyses

OpenBiome donor samples (BioProject: PRJNA544527, subset to whole-genome shotgun samples) were downloaded and processed as described above for metagenomic sequencing processing and mapping to AGORA reference with paired GEMs. Community-scale metabolic models were reconstructed using MICOM for each OpenBiome donor and community metabolic flux profiles were generated as previously described. For each donor taxon, metabolic independence scores were computed using Anvio v8.0, metabolic distance was calculated for each donor taxa relative to each Goll et al. recipient community at baseline, and donor and recipient taxon abundances at T0 were extracted from their respective AGORA-mapped count tables. Shannon diversity scores were computed for all OpenBiome donor microbiomes and all Goll et al. recipient microbiomes at T0, and donor-to-recipient diversity ratios were derived as described above. The random forest classifier trained on the Goll et al. human FMT dataset was then applied to all OpenBiome donor– recipient pairs to predict post-FMT colonization outcomes, enabling *in silico* screening of donor-recipient compatibility across the full OpenBiome cohort.

### Evaluation of Metabolic Competition and Crossfeeding

Species-level pairwise interaction estimates were obtained using MICOM’s summarize_interactions function, which classifies each metabolic exchange as co-consumed (competition), provided, or received (cross-feeding). For each sample, we summed total flux across all co-consumed interactions (competition) and across all provided and received interactions (cross-feeding). We then calculated the net interaction balance (rho) as the normalized difference between total cross-feeding flux and total competition flux: rho = (total cross-feeding flux - total competition flux) / (total cross-feeding flux + total competition flux). This metric ranges from -1 (entirely competition-dominated) to +1 (entirely cross-feeding-dominated) and follows the formulation of Corral Lopez et al^30^.

### Statistical analysis, software, and reproducibility

Data handling was performed with Python (3.9.0) using pandas (v2.3.3) and numpy to generate csv files for downstream statistical analyses and visualizations in R (v4.5.1) using ggplot2 (v4.0.2) and geom_signif extension. Beta diversity analysis was performed with skbio (v0.7.0) in Python. Mahalanobis distance was calculated using scikit-learn (v1.5.1). Significance in metabolic production rates between donors and recipients at T0, engraftment rates across all donor-recipient combinations, and changes in total SCFA production rates between responders and non-responders relative to their T0 baseline were determined by two-sided Mann-Whitney U tests. For multi-group comparisons, p-values were adjusted using the false discovery rate method. Reproducible workflows were implemented through Jupyter Notebooks and R scripts and are available at https://github.com/urtecholabucsd/FMT.

## Availability of Data and Materials

The datasets generated and/or analysed during the current study are available at https://github.com/zetianzhang1/FMT. Raw data for OpenBiome samples were downloaded from BioProject: PRJNA544527. Raw data for the Goll et al. 2020 FMT trial were downloaded from the European Nucleotide Archive (ENA) under the study accession number PRJEB36140. Raw data for murine FMT analysis available through NIH Bioproject: PRJNA1028308.

## ACKNOWLEDGEMENTS

G.U. acknowledges support from the Howard Hughes Medical Institute (GT17584), the NIDDK-funded San Diego Digestive Diseases Research Center (P30 DK120515), the NIH Common Fund’s Faculty Institutional Recruitment for Sustainable Transformation (FIRST) program (NCI U54CA272220), and the NCCIH (R21AT013617). P.D. acknowledges support from NIGMS (R35GM142547).

## Use of Generative AI

The generative AI tool Claude (Opus 4.6, Anthropic) was used during the preparation of this manuscript for coding assistance, editorial proofreading, identification of internal inconsistencies between text and figures, and copy editing of the final draft. All scientific content, analysis, interpretation, and writing were performed by the authors. The authors reviewed and take full responsibility for the final manuscript.

