## Supplementary Figures for "Metagenome-scale Modeling to Assess Microbiome Metabolic Complementarity for Precision Microbiota Transplantation Therapies"

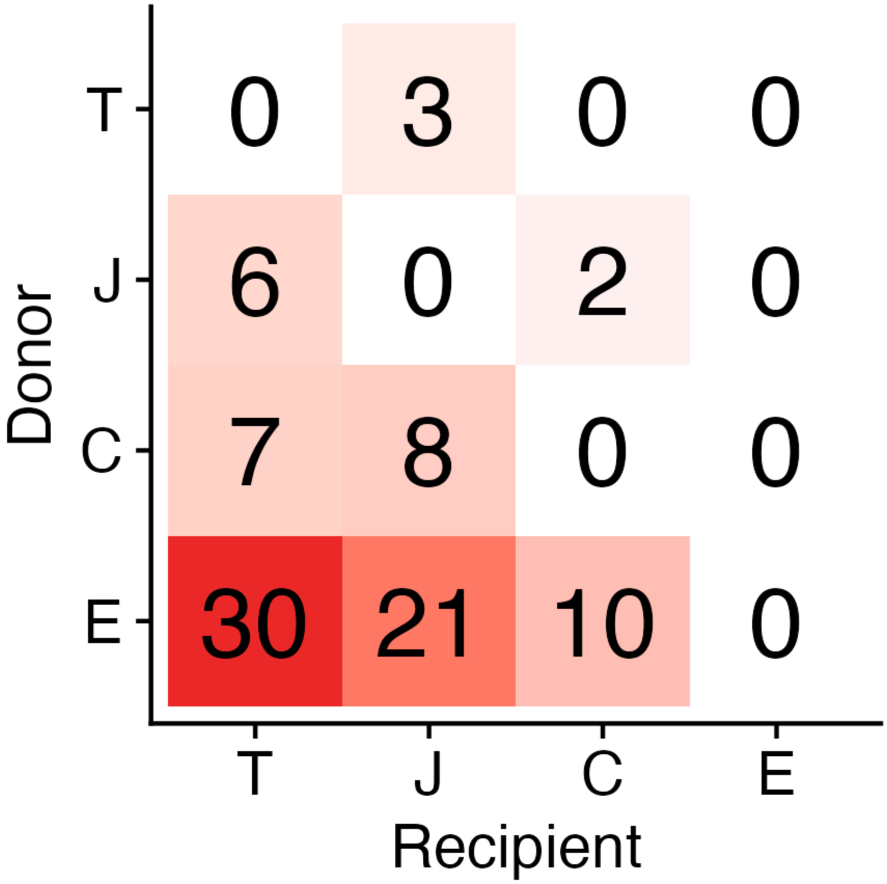


**Figure S1. Variance in the number of taxa transferred via FMT.** Heatmap showing the number of donor taxa that successfully colonized each recipient community across all pairwise FMT combinations between four vendor-distinct murine microbiomes. Rows represent donor communities and columns represent recipients. Cell values indicate the count of transferred taxa. Envigo microbiota exhibited the highest colonization capacity across recipients while remaining fully resistant to reciprocal colonization. Conversely, Taconic communities were the most receptive to incoming donor taxa but engrafted poorly when used as donors.

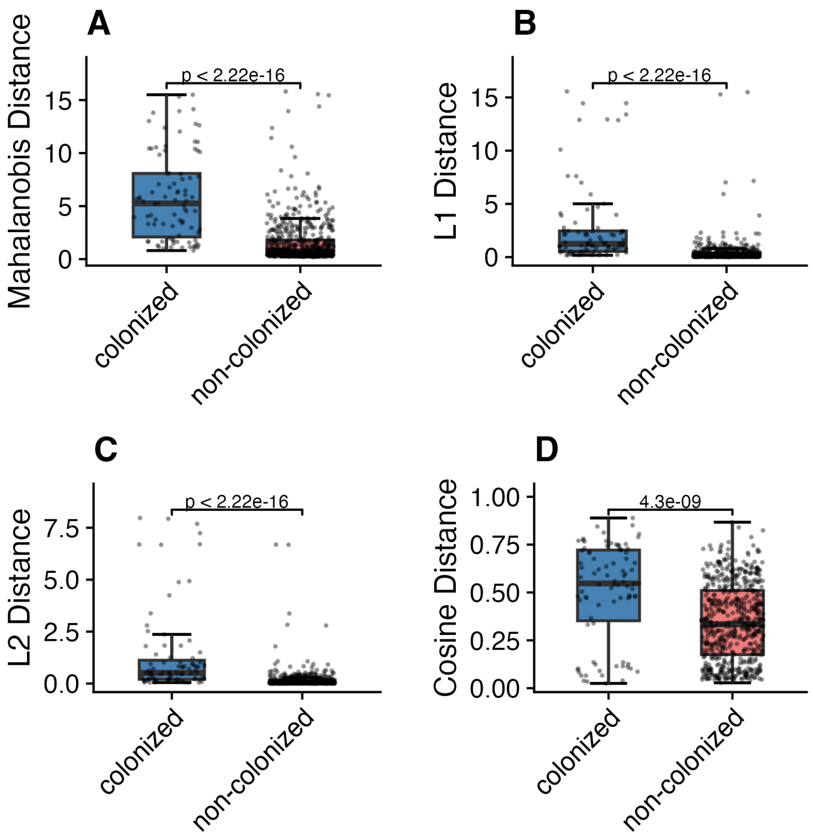


**Figure S2. Comparison of distance metrics for discriminating colonizing from non-colonizing taxa.** Boxplots show the distribution of (**A**) Mahalanobis distance, (**B**) L1 distance, (**C**) L2 distance, and (**D**) cosine distance between colonizing and non-colonizing donor taxa across all pairwise vendor FMT cohorts. All four metrics significantly distinguished colonizers from non-colonizers (Wilcoxon rank-sum test), but Mahalanobis distance yielded the greatest separation between groups. Boxes span the interquartile range with the median indicated by the horizontal line; whiskers extend to 1.5x the interquartile range. Individual data points are overlaid with jitter.

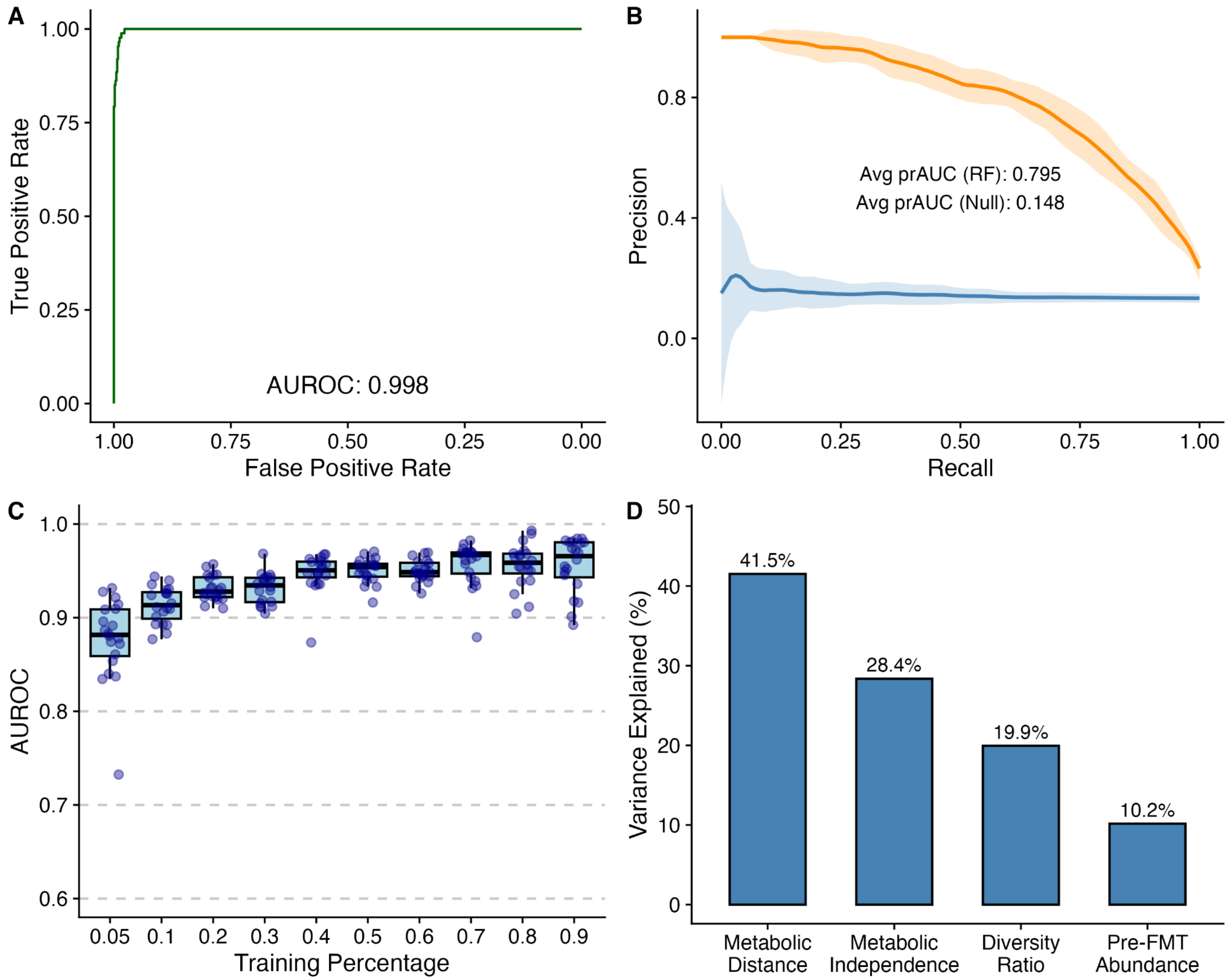


**Figure S3. Random forest classifier accurately predicts colonization outcomes. A)** Receiver operating characteristic (ROC) curve for the random forest classifier trained on metabolic distance, metabolic independence, pre-FMT donor taxon abundance (log10-transformed), and donor-to-recipient Shannon diversity ratio. **B)** Precision-recall curve for the same classifier evaluated over 20 iterations of Monte Carlo cross-validation (50/50 train/test split). Orange line and shading indicate the mean and standard deviation of precision across iterations for the random forest model; blue line and shading indicate a null model generated by label permutation. Average precision-recall AUC (prAUC) values are annotated for both models. **C)** Classifier robustness across training set sizes. Boxplots show AUROC distributions from 20 iterations of Monte Carlo cross-validation at each training proportion (5% to 90%). Individual iterations are overlaid as points. **D)** Relative contribution of each predictor to colonization classification, assessed by likelihood-ratio chi-squared test (Type II ANOVA) on a logistic regression model with the same predictors. Bars show the percentage of total variance explained by each feature.

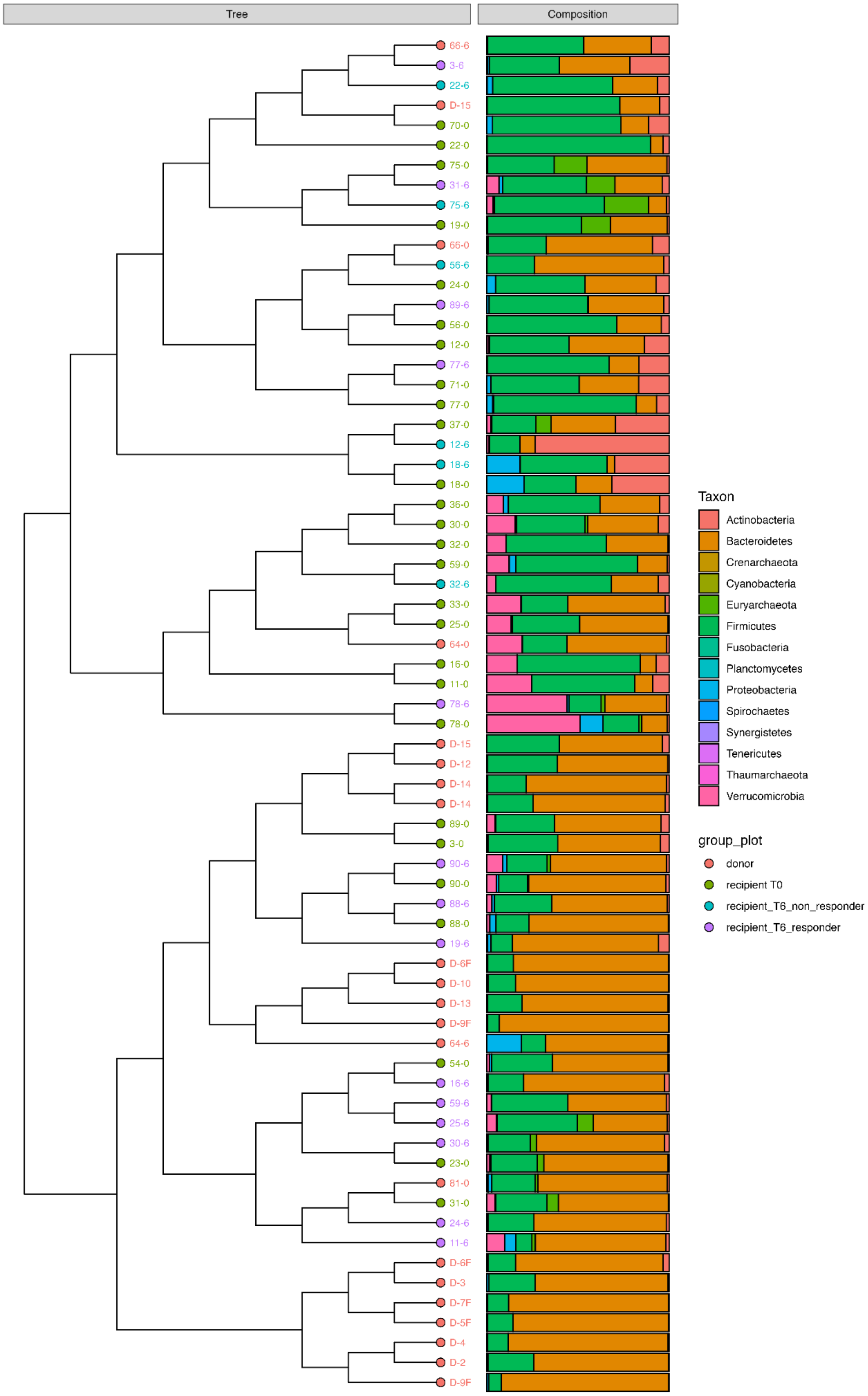


**Figure S4. Hierarchical clustering of taxonomic composition across donors, recipients, and post-FMT timepoints.** Dendrogram (left) shows hierarchical clustering of Bray-Curtis dissimilarities between all samples. Stacked bar plots (right) display phylum-level relative abundance for each sample. Tip labels are colored by sample group: donors (red), recipients at baseline (green, T0), post-FMT responders at six months (purple, T6), and post-FMT non-responders at six months (cyan, T6). Responders were defined as patients achieving a 75-point or greater decrease in IBS Severity Scoring System (IBS-SSS) at three months. Donor samples clustered together and were characterized by higher relative abundance of *Bacteroidales* compared to recipient samples at baseline. Sample identifiers denote patient number and timepoint (e.g., 12-0 = patient 12 at T0; 12-6 = patient 12 at T6; D-3 = donor 3).
